# Diverse noncanonical PAMs recognized by SpCas9 in human cells

**DOI:** 10.1101/671503

**Authors:** Ziying Hu, Daqi Wang, Chengdong Zhang, Shuai Wang, Siqi Gao, Linghui Hou, Hongyan Wang, Yongming Wang

## Abstract

The CRISPR/Cas9 system derived from *Streptococcus pyogenes* (SpCas9) provides unprecedented genome editing capabilities, but the potential for off-target mutations limits its application. In addition to NGG protospacer adjacent motif (PAM), off-target mutations are also associated with noncanonical PAMs, which have not yet been systematically evaluated. Here, we developed a highly sensitive approach that allows systematically analyzing PAM sequences in human cells, and identified multiple alternative PAMs recognized by SpCas9.

## INTRODUCTION

CRISPR/Cas9 system has a broad range of research and medical applications. This system consists of a Cas9 nuclease and a guide RNA (gRNA), which form a Cas9-gRNA complex, recognizing a target sequence with a downstream PAM and induce a site-specific double-strand break (DSB) ^1–3^. Although researchers have repurposed many different CRISPR/Cas systems for genome targeting, SpCas9 is the most extensively studied and applied system to date due to its high efficiency and simple PAM requirement ^1–6^. In addition to NGG PAM, several noncanonical PAMs including NAG, NCG, NAG and NNGG recognized by SpCas9 have been identified ^7, 8^. Noncanonical PAMs are associated with low efficiency of on-target activity, but they must be considered as potential off-target sites ^9, 10^. Therefore, it is crucial to systematically investigate the noncanonical PAMs associated with *Sp*Cas9.

The essential of the PAM has spurred the development of multiple approaches to identify PAMs, including in silico PAM prediction ^11, 12^, in vivo PAM library screening in bacteria ^8, 13^ and in vitro PAM library screening ^14–16^. In silico PAM prediction remains limited by the availability of matching phage or plasmid DNA sequences in genomic databases, and obtained hits may include mutated escape-PAMs. In vivo PAM library screening relies on negative screens (plasmid depletion experiments) in bacteria, which gives high background. In vitro PAM library screening can be highly sensitive to the assay conditions ^15^. These shortcomings highlight the need for screens that overcome these limitations. Here, we developed a GFP reporter-based in vivo PAM library screening assay with two distinct features: i) the assay is performed in human cellular context; ii) the assay is sensitive enough to detect single cleavage event.

## RESULTS AND DISCUSSION

In this PAM library screening system, a protospacer containing downstream randomized DNA sequences is inserted between ATG start codon and GFP coding sequence, disrupting GFP expression by frameshift mutation (Figure 1a). When CRISPR/Cas9 cleaves the protospacer and generates small insertions/deletions (indels), a portion of cells will restore the GFP reading frame. The reporter system was delivered into HEK293T cells by lentiviral infection to get stable cell line. Two days after transfection of SpCas9 with corresponding gRNA, GFP-positive cells could be observed (Figure 1b). In contrast, transfection of SpCas9 alone could not induce GFP expression (Figure 1b). The GFP-positive cells were sorted out, and the protospacer with randomized DNA sequences were PCR-amplified for deep sequencing. Only in-frame mutations were considered as novel mutations induced by SpCas9. Since indels could disrupt the randomized DNA sequences, only GCG triple-nucleotide followed by 7-bp sequences in front of GFP coding sequence was used for PAM analysis. Deep sequencing analysis revealed that indels associated with different PAMs could be detected by this assay (Figure 1c). Both sequence logo and PAM wheel captured the canonical NGG as the most enriched PAM, but the sequences other than NGG can also be observed (Figure 1d-e).

**Figure 1.**
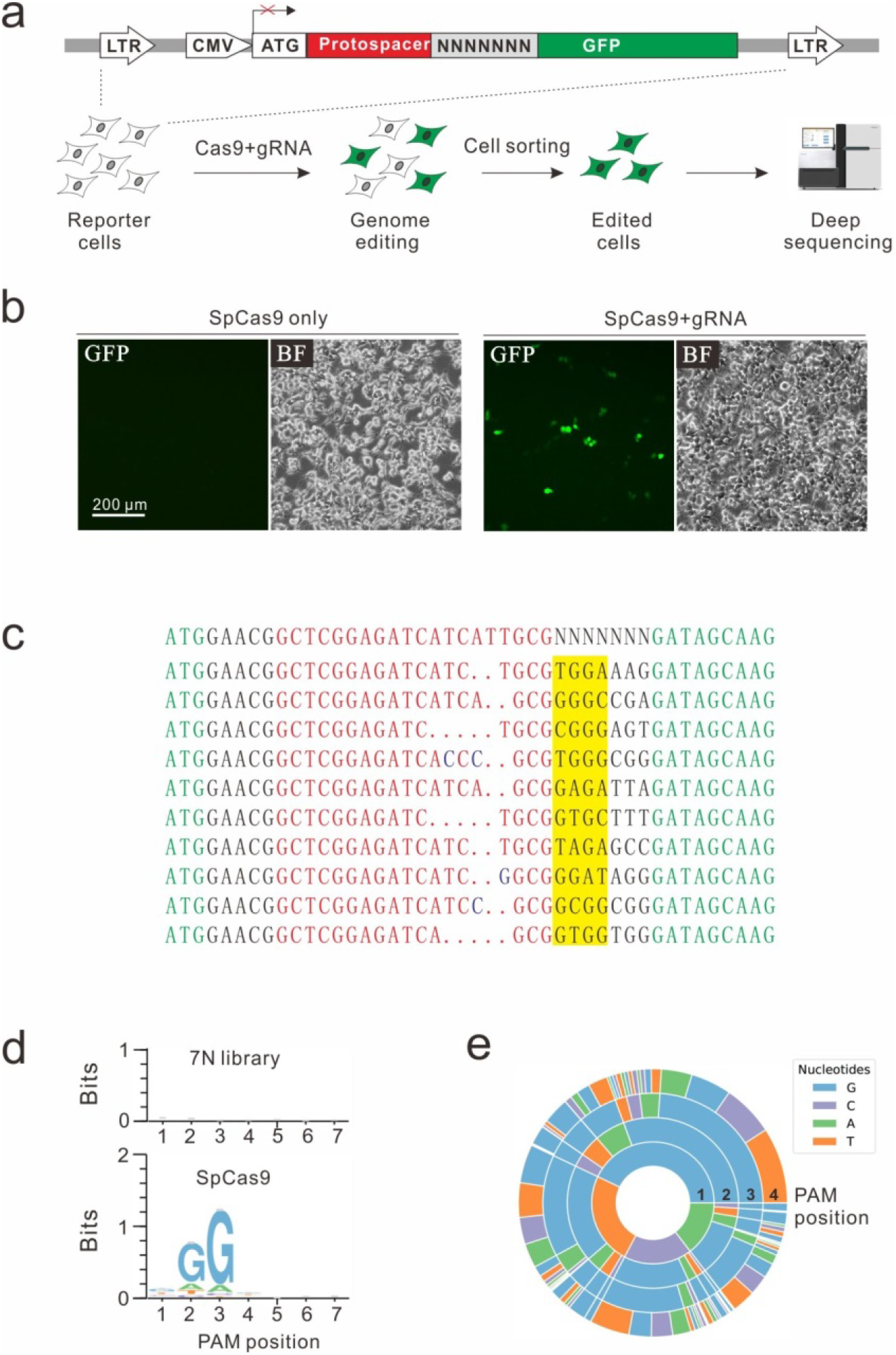
A GFP reporter assay for PAM screening. a. Schematic diagram of the GFP reporter assay. A lentiviral vector contains a CMV-driven GFP. A target sequence followed by 7-bp random sequence was inserted between ATG and GFP coding sequence, disrupting GFP expression. The library was stably integrated into HEK293T cells. After genome editing, a portion of cells will restore GFP expression. GFP-positive cells are sorted out and PAM sequences were PCR-amplified for deep-sequencing analysis. b. Transfection of SpCas9 and gRNA resulted in GFP expression, while transfection of SpCas9 alone could not induce GFP expression. c. Deep-sequencing revealed that targets with multiple PAMs could be edited. GFP sequence is shown in green; target sequence is shown in red; insertion mutations are shown in blue; 4-bp PAM sequences are highlight in yellow. d. Sequence logo was generated from PAM screening assay. e. PAM wheel was generated from PAM screening assay.

Since nucleotide bias can be observed from position 1 to 4 in the sequence logo (Figure 1d), we analyzed the efficiency ratio for all possible NNNN PAM sequences. 138 out of 256 PAMs can be edited (Suppl. Table 1). The top 74 PAMs included all NNGG, NAGN, NGAN and GGYN PAMs with efficiency ratio over 1‰. The remaining 64 PAMs only displayed minimal activity. The most efficient PAM was NGGN, as expected, followed by NNGG, NAGN, NGAN and GGYN (Figure 2a and Suppl. Table 1). Interestingly, some noncanonical PAMs such as GTCC, GCGG and GAGT displayed similar efficiency to NGGN PAMs (Figure 2a).

**Figure 2.**
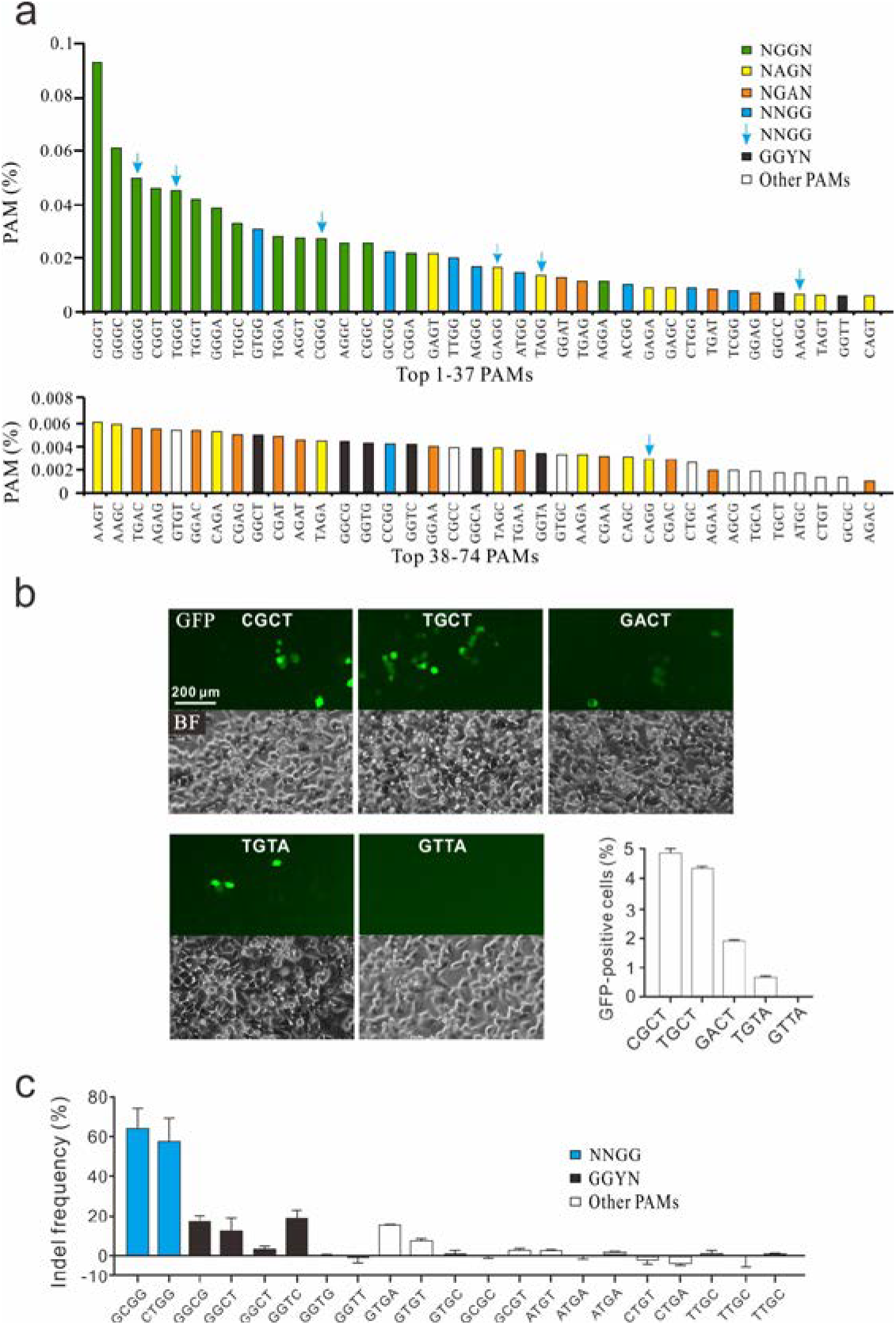
PAM sequence analysis. a. Efficiency ratio of top 74 PAMs. PAM sequences were shown in tetramer. NNGG PAM overlapped with other PAMs was indicated by arrows. b. Five GFP reporter constructs were isolated from PAM library and tested for genome editing. c. Indel frequency of endogenous target sequences associated with noncanonical PAMs.

To confirm PAMs identified in the library, we isolated five GFP reporter constructs with different PAM sequences from the PAM library and established stable cell lines for each construct. Transfection of SpCas9 and gRNA induced GFP expression for four PAMs (CGCT, TGCT, GACT and TGTA), indicating that cleavage occurred with these PAMs (Figure 2b). GFP-positive cells were not detectable for GTTA PAM, consistent with the PAM screening results that GTTA PAM had no activity. We further tested 21 endogenous targets with noncanonical PAMs. Two NNGG PAMs displayed very high activity, with indel rates of 64.3% and 57.6% for GCGG and CTGG, respectively (Figure 2c). Four GGYN PAMs (GGTC, GGCG, GGCT and GGCT) as well as GTGA and GTGT PAMs also displayed significant activity. Other PAMs displayed minimal activity.

In summary, we developed a highly sensitive GFP reporter-based method for PAM screening and identified multiple novel noncanonical PAMs for SpCas9. When researchers design a gRNA, target sequences associated with these noncanonical PAMs should be considered as potential off-target sequences.

## MATERIALS AND METHODS

### Cell culture and transfection

HEK293T cells were maintained in Dulbecco’s Modified Eagle Medium (DMEM) supplemented with 10% FBS (Gibco) and 1% antibiotics at 37 °C with 5% CO_2_. For PAM library screen, HEK293T cells were plated into 10cm dishes and transfected at ~ 60% confluency with SpCas9-gRNA-expressing plasmid (15 μg) using Lipofectamine 2000 (Life Technologies). For PAM validation at endogenous site, HEK293T cells were seeded on 48-well plates and transfected with SpCas9-gRNA plasmid (500 ng) using Lipofectamine 2000.

### The PAM library construction

The DNA oligonucleotides consisted of two side sequences for Gibson Assembly, 25-bp targeting sequence and seven random bases adjacent to the target (**Table 1**). The DNA oligonucleotides were synthesized and purchased from GENEWIZ. Full-length oligonucleotides were PCR-amplified using Q5 High-Fidelity 2X Master Mix (NEB), size-selected using a 3% agarose EGel EX (Life Technologies, Qiagen), and purified using MinElute Gel Extraction Kit (Qiagen). PCR products were cloned into Lentiviral vector by Gibson Assembly (NEB) and purified with Agencourt AMPure XP SPRI beads (Beckman Coulter). The Gibson Assembly products were electroporated into MegaX DH10B^TM^ T1^R^ Electrocomp^TM^ Cells (Invitrogen) using a GenePulser (BioRad). The bacterial were added into recovery media and grown at 32 °C, 225 rpm for 14 h. The plasmid DNA was extracted from bacterial cells using Endotoxin-Free Plasmid Maxiprep (Qiagen).

### Lentivirus production

For PAM library packaging, HEK293T cells were seeded in three 10 cm dishes and transfected at ~40% confluency. For each dish, 12 μg of PAM plasmid, 9 μg of psPAX2, and 3 μg of pMD2.G were transfected with 60 μl of Lipofectamine 2000 (Life Technologies). Virus was harvested twice at 48 h and 72 h post-transfection. The virus was concentrated using PEG8000 (no. LV810A-1, SBI, Palo Alto, CA), dissolved in PBS and stored at −80 °C. For single PAM packaging, HEK293T cells were seeded into 6-well plates, 1.2 μg of PAM plasmid, 0.9 μg of psPAX2, and 0.3 μg of pMD2.G were transfected with 5 μl of Lipofectamine 2000. Virus was harvested twice at 60 h post-transfection.

### PAM library screening experiments

HEK293T cells were plated into 15 cm dish at ~30% confluence. After 24 h, cells were infected with PAM library lentivirus with at least 1000-fold coverage of each PAM. 24 h after infection, the cells were selected with 2 µg/ml of puromycin for 5 days. The GFP negative cells were sorted with a MoFlo XDP machine (Beckman Coulter) and seeded into 10 cm dishes. Then the PAM library cells were transfected with Cas9-gRNA expressing plasmid. Three days after editing, the GFP positive cells were sorted out by MoFlo XDP machine and the genomic DNA was isolated using TIANamp Genomic DNA Kit (TIANGEN) following the manufacturer’s instructions. PCR fragments for deep-sequencing were generated in two-step PCR reactions. First, the target region was PCR-amplified using primers Deep-F1/R1 with 25 cycles using Q5 High-Fidelity 2X Master Mix (NEB). Second, 3 μl of PCR products from first step was amplified by primer P5-adapter-F and P7-adapter-R for 15 cycles (Primer sequence in **Table 1**). The PCR products were purified using Gel Extraction Kit (Qiagen) and were sequenced on Illumina HiSeq X by 150-bp paired-end sequencing.

### PAM sequence analysis

Twenty base pair sequences flanking the target sequence (GAACGGCTCGGAGATCATCATTGCGNNNNNNN) was used to fix the target sequence. GCG followed 7-bp random sequence and GFP coding sequence was used to fix the PAM sequence. Target sequences with in-frame mutations and contact 7-bp random sequence were used for PAM analysis. Then the sequence composition of the last 7bp base was analyzed.

### Verification of individual PAM sequence with GFP reporter constructs

Five plasmids containing different PAM sequences were selected from the PAM library. After packing, the lentivirus containing single PAM sequence were added to the 12-well HEK293T cells. After 7 days of puromycin screening, the GFP negative cells were sorted by MoFlo XDP machine. The sorted cells were seeded into 24-well and transfected with SpCas9-gRNA plasmid (800ng) by Lipofectamine 2000. Five days after editing, the positive cells were analyzed on the Calibur instrument (BD). Data were analyzed using FlowJo.

### Test of PAM activity at endogenous site

In order to test the noncanonical PAM activity, we selected 21 endogenous sites with different PAMs for genome editing (**Table 1**). HEK293T cells were transfected with SpCas9 and gRNA expressing plasmid, and the cells were screened with 2 µg/ml of puromycin for 6 days. The genomic DNA was isolated, the target sites were amplified with two-step PCR. Indel frequency was detected by deep-sequencing analysis.

## REFERENCES

1. Jinek, M. et al. A programmable dual-RNA-guided DNA endonuclease in adaptive bacterial immunity. Science 337, 816–821 (2012).

2. Cong, L. et al. Multiplex genome engineering using CRISPR/Cas systems. Science 339, 819–823 (2013).

3. Mali, P. et al. RNA-guided human genome engineering via Cas9. Science 339, 823–826 (2013).

4. Wang, H.Y. et al. One-Step Generation of Mice Carrying Mutations in Multiple Genes by CRISPR/Cas-Mediated Genome Engineering. Cell 153, 910–918 (2013).

5. Fu, Y., Sander, J.D., Reyon, D., Cascio, V.M. & Joung, J.K. Improving CRISPR-Cas nuclease specificity using truncated guide RNAs.

6. Xie, Y. et al. An episomal vector-based CRISPR/Cas9 system for highly efficient gene knockout in human pluripotent stem cells. Scientific reports 7, 2320 (2017).

7. Doench, J.G. et al. Optimized sgRNA design to maximize activity and minimize off-target effects of CRISPR-Cas9. Nature biotechnology 34, 184–191 (2016).

8. Jiang, W., Bikard, D., Cox, D., Zhang, F. & Marraffini, L.A. RNA-guided editing of bacterial genomes using CRISPR-Cas systems. Nature biotechnology 31, 233–239 (2013).

9. Lin, Y. et al. CRISPR/Cas9 systems have off-target activity with insertions or deletions between target DNA and guide RNA sequences. Nucleic acids research 42, 7473–7485 (2014).

10. Li, J. et al. Whole genome sequencing reveals rare off-target mutations and considerable inherent genetic or/and somaclonal variations in CRISPR/Cas9-edited cotton plants. Plant biotechnology journal 17, 858–868 (2019).

11. Horvath, P. et al. Diversity, activity, and evolution of CRISPR loci in Streptococcus thermophilus. Journal of bacteriology 190, 1401–1412 (2008).

12. Mojica, F.J., Diez-Villasenor, C., Garcia-Martinez, J. & Almendros, C. Short motif sequences determine the targets of the prokaryotic CRISPR defence system. Microbiology 155, 733–740 (2009).

13. Esvelt, K.M. et al. Orthogonal Cas9 proteins for RNA-guided gene regulation and editing. Nat Methods 10, 1116–1121 (2013).

14. Ran, F.A. et al. In vivo genome editing using Staphylococcus aureus Cas9. Nature 520, 186–191 (2015).

15. Karvelis, T. et al. Rapid characterization of CRISPR-Cas9 protospacer adjacent motif sequence elements. Genome biology 16, 253 (2015).

16. Pattanayak, V. et al. High-throughput profiling of off-target DNA cleavage reveals RNA-programmed Cas9 nuclease specificity. Nature biotechnology 31, 839–843 (2013).

